# Multimodal acoustic-electric trigeminal nerve stimulation modulates conscious perception

**DOI:** 10.1101/2023.03.21.533632

**Authors:** Min Wu, Ryszard Auksztulewicz, Lars Riecke

## Abstract

Multimodal stimulation has the potential to reverse pathological neural activity and alleviate symptoms in neuropsychiatric diseases. However, the reliability of this approach and the mechanisms through which it improves consciousness remain largely unknown. We investigated the effects of multimodal stimulation combining music stimulation with electrical trigeminal nerve stimulation in healthy human participants. We assessed conscious perception before and after acoustic-electric stimulation and investigated the mechanisms underlying the putative stimulation effects. Our results show that (1) acoustic-electric stimulation improves conscious tactile perception in healthy human participants without a concomitant change in auditory perception, (2) this improvement is caused by the interplay of the acoustic and electric stimulation rather than any of the unimodal stimulation alone, and (3) the effect of acoustic-electric stimulation on conscious perception correlates with inter-regional connection changes in a recurrent neural processing model. These findings provide evidence that multimodal acoustic-electric stimulation can promote conscious perception and offer insights into its underlying mechanisms.

## Introduction

Consciousness is an important and ubiquitous aspect of our daily lives. Many pathologic brain states, such as epileptic seizures, stroke and encephalitis, can lead to impaired consciousness. In particular during the COVID-19 pandemic, up to 19.6% of patients with COVID-19 suffered from impaired consciousness (Mao et al., 2020; Romero-Sánchez et al., 2020). Consciousness impairment can have devastating consequences for safety and quality of life; therefore, it is particularly important to develop effective approaches to promote consciousness.

Over the past decades, an increasing variety of consciousness-promoting therapies have been proposed. These therapies include pharmacological treatments (e.g., zolpidem, apomorphine) (Sanz et al., 2019; Thonnard et al., 2013) and nonpharmacologic interventions, such as sensory or electric stimulation (Cooper et al., 2005; Thibaut et al., 2019). A recent approach, inspired by the increasing knowledge of the neural mechanisms underlying consciousness, involves non-invasive electric stimulation of the trigeminal nerve (Fan et al., 2019; Wu et al., 2022). This cranial nerve projects facial sensation to the brainstem and the cortex, and connects to the reticular activating system, thalamus, insula, and somatosensory brain structures (Simpson et al., 1997)—regions which have been proposed to play a role in consciousness (Gallace and Spence, 2010; Schiff et al., 2007; Steriade, 1996).

Trigeminal nerve stimulation (TNS) has already been widely used in many neurological disorders (DeGiorgio et al., 2013; McGough et al., 2019) and first applications of it in consciousness research have yielded promising results. It has been observed that TNS of rats with impaired consciousness can upregulate neuropeptide hypocretin in the lateral hypothalamus and this activation can promote consciousness recovery (Zheng et al., 2021). In other animal models with traumatic brain injury, TNS has been demonstrated to enhance cerebral blood flow, protect the blood-brain barrier, and reduce brain edema (Chiluwal et al., 2017; Yang et al., 2022). Similar effects of TNS have been shown in human patients diagnosed with disorders of consciousness (DOC). In a single-case study, Fan et al. reported an improvement of consciousness after TNS in a DOC patient (Fan et al., 2019). Similarly, Dong et al. reported consciousness benefits after four weeks of TNS in eight out of 21 DOC patients (Dong et al., 2022). Therefore, both animal and human findings suggest that TNS may be an effective approach for promoting the recovery of consciousness.

Compared to unimodal stimulation, multimodal stimulation has been shown to modulate more widespread brain regions and cause stronger neural activation within the multisensory regions (Godenzini et al., 2021; Markovitz et al., 2015; Marks et al., 2018). For example, combined acoustic and visual stimulation in Alzheimer’s disease (AD) mice has been found to reduce amyloid load (a pathological hallmark of AD) across much broader cortical regions than acoustic or visual stimulation alone (Martorell et al., 2019). Similar benefits of multimodal stimulation have been observed in tinnitus studies, which found that combined acoustic and electric stimulation, but not unimodal stimulation, can reverse tinnitus-related pathological neural activity and alleviate tinnitus symptoms (Engineer et al., 2011; Marks et al., 2018). These findings suggest that multimodal stimulation allows for a more effective treatment of some cognitive disorders than unimodal stimulation.

Inspired by these findings, we recently conducted a stimulation study in which we combined rhythmic acoustic music stimulation and rhythmic transcutaneous electrical TNS in DOC patients (Wu et al., 2022). We found that multimodal acoustic-electric stimulation in the gamma band (40Hz) can promote both gamma neural activity and re-emergence of consciousness. Given the evidence above, multimodal acoustic-electric stimulation appears to be a promising approach for promoting consciousness. As the approach is still novel, its reliability and functional principle are still poorly understood and require further investigation.

In the aforementioned DOC patient studies, the level of consciousness was assessed by clinicians with a questionnaire, the Coma-Recover Scale Revised (CRS-R). Although this scale currently constitutes the gold standard for consciousness assessment (Seel et al., 2010), it highly depends on the assessor’s experience and can lead to a high rate of misdiagnosis in patients with motor impairment (Schnakers et al., 2009). An alternative measure that is commonly used in studies of neural correlates of consciousness exploits participants’ subjective experience of sensory input, as quantified with perceptual performance on a near-threshold target-detection task (Eklund and Wiens, 2019). In this assessment, healthy participants are asked to report whether they are aware or unaware of a sensory stimulus that is repeatedly presented at a fixed intensity near the perceptual detection threshold. While this measure can serve merely as a proxy for consciousness, it has the advantage that it directly reflects the participant’s subjective experience and can be readily coupled with objective measures. For example, neural markers of conscious perception can be identified by comparing neural responses to detected vs. undetected identical stimuli while controlling for confounding variations in sensory input. Results of previous EEG studies using this approach suggest that conscious perception involves two prominent event-related potential (ERP) components: an early negative component in sensory regions 120-200 ms after the stimulus onset and a late positive component in occipital-parietal regions 250-500 ms after the stimulus onset (Dembski et al., 2021; Eklund and Wiens, 2019; Koivisto and Revonsuo, 2010). If multimodal acoustic-electric stimulation can reliably promote consciousness, one would expect it to effectively enhance these alternative measures of consciousness.

Despite the converging results of the aforementioned AD and tinnitus studies, the empirical evidence for stronger benefits from multimodal (compared with unimodal) stimulation for consciousness is still very limited. So far, consciousness benefits from multimodal stimulation have been investigated primarily by stimulating in different sensory modalities consecutively (Cheng et al., 2018; Megha et al., 2013), rather than simultaneously (Wu et al., 2022). Thus, it is still unclear whether the consciousness benefit from multimodal acoustic-electric stimulation results from the multimodal nature of the stimulation. More generally, the mechanism by which the multimodal stimulation may improve consciousness is still unclear. Therefore, it remains to be determined whether and how consciousness benefits from multimodal acoustic-electric stimulation are driven by the acoustic or electric stimulation, or their simultaneous combination.

The present study aimed to investigate (1) whether the effect of multimodal stimulation combining music stimulation and electrical TNS (hereafter shortly called “acoustic-electric stimulation”) on consciousness is reproducible in healthy participants using behavioral and neural measures of conscious perception in a target-detection task; (2) whether the effect is primarily driven by the combination of the acoustic and electric stimulation or any of the unimodal inputs alone; and (3) potential neural mechanisms underlying the putative effects of acoustic-electric stimulation. To achieve aims 1 and 2, we performed two experiments in healthy human participants using a double-blinded, randomized, crossover design. We applied acoustic-electric and acoustic-only stimulation (Experiment 1, Fig.1A), and electric-only and electric-sham stimulation (Experiment 2, Fig.1B). We assessed conscious perception before and after the stimulation based on participants’ perceptual performance on tactile and auditory target-detection tasks and electroencephalography (EEG) responses to undetected vs. detected targets (Fig.2). To achieve aim 3, we applied a dynamic-causal modeling (DCM) approach to fit a biologically plausible neural network model to the EEG data. DCM is a useful tool to study the neural architecture underlying observed electrophysiological features in terms of effective connectivity (Stephan et al., 2010). Previous EEG studies have utilized this approach to investigate neural mechanisms of tactile conscious perception and found a potential role of recurrent neural processing in the cortex (Auksztulewicz and Blankenburg, 2013; Auksztulewicz et al., 2012). Previous consciousness studies have demonstrated that the disruption of connectivity is a candidate mechanism of impaired consciousness, and restoration of brain connectivity is accompanied by recovery of consciousness (Boly et al., 2011; Laureys et al., 2000). On this basis, we hypothesized that changes in effectivity connectivity might be the mechanisms by which acoustic-electric stimulation induces consciousness benefits. We analyzed the connectivity parameters of the best-fitting neural network model to test whether acoustic-electric stimulation can modulate estimates of inter-regional cortical connections.

**Figure 1.**
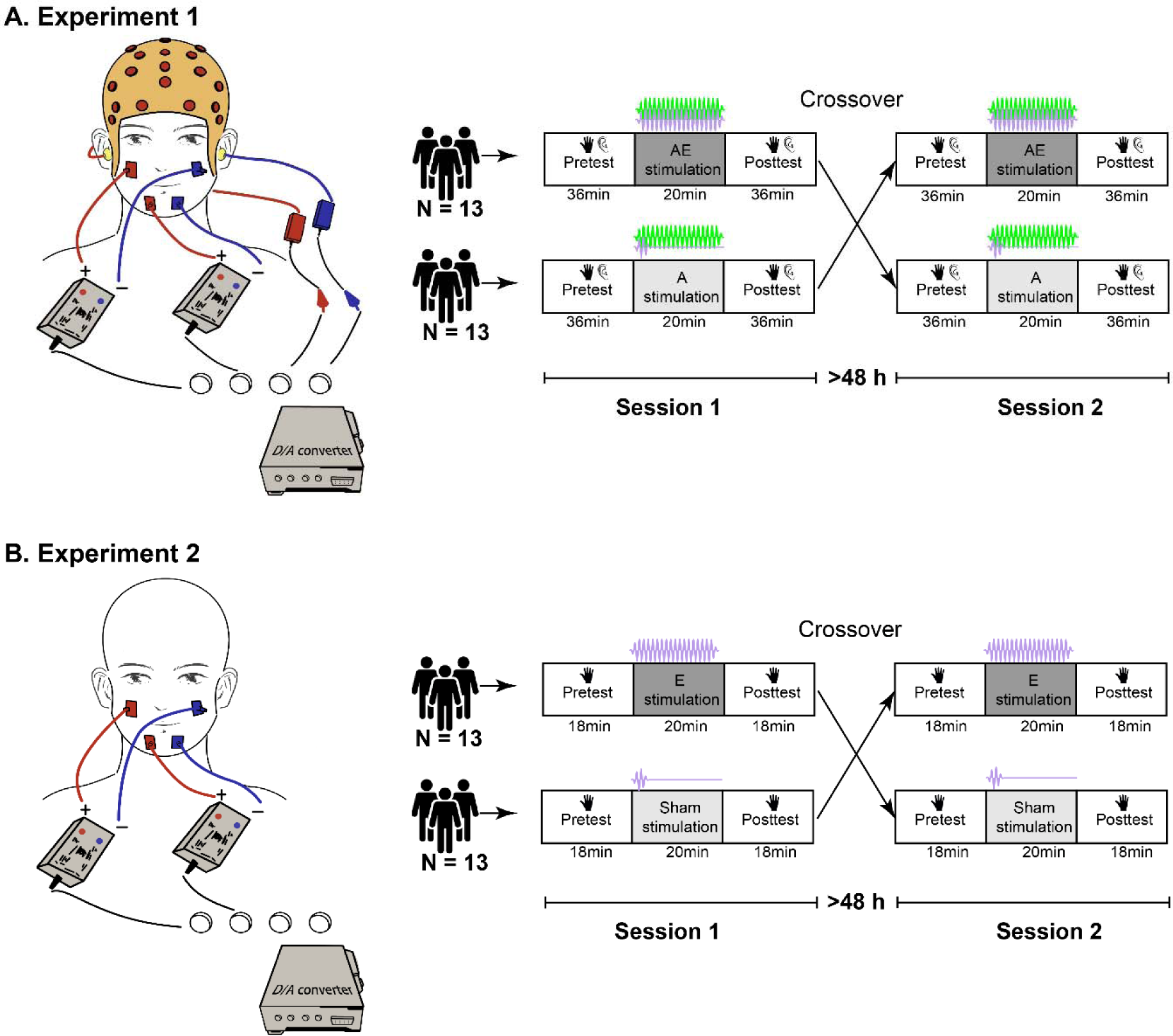
Stimulation approach and study design. **(A).** Experiment 1: Multimodal acoustic-electric TN stimulation involved the simultaneous application of music via earphones and electric current via electrodes attached to the middle and lower parts of the participant’s face (see red and blue squares). The crossover design involved two sessions separated by more than 48 hours. Each session consists of three phases: assessment of conscious perception before stimulation (pretest), application of acoustic-electric/acoustic-only stimulation, and assessment of conscious perception after stimulation (posttest). **(B).** Experiment 2: same as (A), but the stimulation involved electric-only and electric-sham stimulation and the assessment focused only on conscious tactile perception.

**Figure 2.**
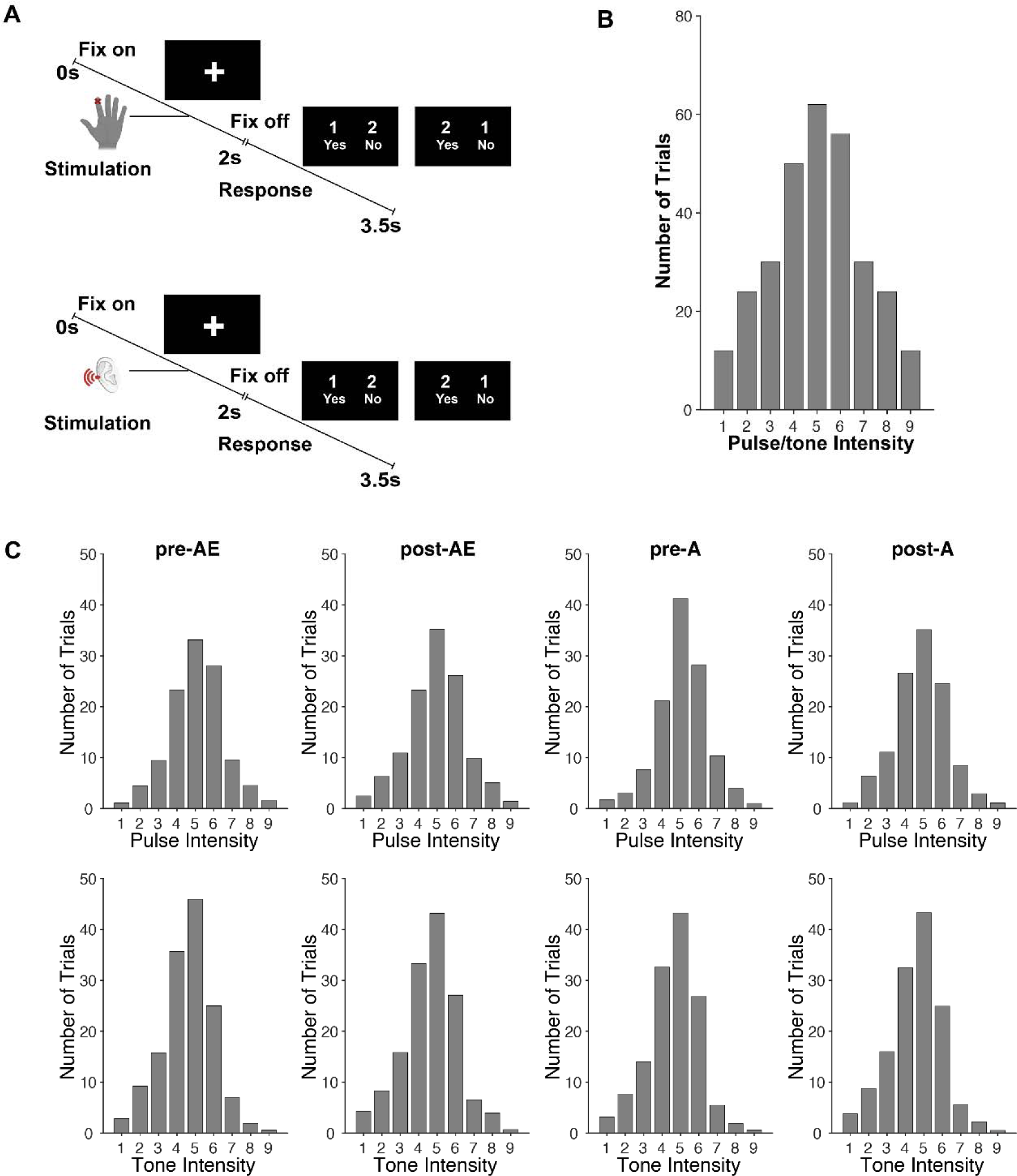
Trial design and trial distributions. **(A).** Schematic of an exemplary tactile-detection trial (upper) and an auditory-detection trial (bottom) presented in different blocks. A trial started with the presentation of a fixation cross on the screen. A tactile or auditory stimulus followed with a random delay 1000-1500 ms after the fixation onset. Next, a 1500-ms response screen was displayed, instructing participants to respond whether they had detected a stimulus or not. **(B).** Intensity distribution of the presented trials. The number of trials across intensity levels followed a normal distribution, with a maximal number of trials at the threshold level (intensity 5). **(C).** Intensity distribution of trials after subsampling for the neural data analysis. The subsampling resulted in a matched number of detected and undetected trials, for each intensity.

We hypothesized that (1) acoustic-electric stimulation elicits an improvement of conscious perception in healthy human participants, as indicated by increases in both detection performance and awareness-related ERP components after vs before the stimulation, (2) this improvement in conscious perception is larger after acoustic-electric stimulation than after acoustic or electric stimulation alone, and (3) the putative effect of acoustic-electric stimulation on consciousness correlates with inter-regional connection changes in a recurrent neural processing model.

## Results

In both experiments, participants’ subjective reports of the received electric stimulation did not differ between the two experimental sessions, which involved respectively verum and sham electric stimulation (*p* < 0.05, Fisher’s exact test). This suggests that participants could not reliably distinguish between the presence and absence of the electric stimulation. No participant reported any side effect. Two participants reported that they were familiar with the music (i.e., the acoustic stimulation) in both sessions (familiarity scores: 3 and 4 out of 5), while all other participants reported they were completely unfamiliar with the music (familiarity score: 0 out of 5).

### Acoustic-electric stimulation but not acoustic-only stimulation modulates conscious tactile perception

To assess whether acoustic-electric stimulation or acoustic-only stimulation can improve subsequent conscious perception, we compared stimulus-detection thresholds (interpreted as response criterion, as e.g. a leftward shift of the psychometric curve represents a bias towards reporting “detected” (Gold and Ding, 2013; Kar and Krekelberg, 2014; Ruzzoli et al., 2010)) and psychometric slopes (interpreted as perceptual sensitivity, as e.g. steeper slope represents higher sensitivity (Gold and Ding, 2013; Parker and Newsome, 1998; Zazio et al., 2019)) on tactile and auditory target-detection tasks after vs before stimulation in Experiment 1 (Fig.1A). We found a significant decrease in detection threshold, but not in slope, in the tactile task after vs before acoustic-electric stimulation (threshold: t_25_ = 3.936, *p* = 5.8 × 10^-4^, effect size d = 0.772; slope: t_25_ = −0.061, *p* = 0.952, d = −0.012, Fig.3A-C). In contrast, we found no significant change in either detection threshold or slope in the tactile task after vs before acoustic-only stimulation (threshold: t_25_ = 1.831, *p* = 0.079, d = 0.359; slope: t_25_ = 1.030, *p* = 0.313, d = 0.202, Fig.3A-C). This observation of a behavioral gain in tactile perception after acoustic-electric stimulation (vs. acoustic-only stimulation) was statistically supported by results from a two-way repeated-measures ANOVA, which revealed a significant *stimulation* (acoustic-electric stimulation versus acoustic-only stimulation) × *time* (pretest versus posttest) interaction (F_1,25_ = 8.572, *p* = 0.007, effect size = 0.255). For conscious auditory perception, we found no significant change in detection threshold or slope after vs before acoustic-electric stimulation (threshold: t_25_ = 0.707, *p* = 0.707, d = 0.075; slope: t_25_ = 0500, *p* = 0.621, d = 0.098) or acoustic-only stimulation (threshold: t_25_ = −1.123, *p* = 0.272, d = −0.220; slope: t_25_ = −0.123, *p* = 0.903, d = −0.024) (Fig.3D-F). These results indicate that (1) acoustic-electric stimulation, but not acoustic-only stimulation, improved subsequent conscious tactile perception, and this improvement affected exclusively participants’ response criterion, not perceptual sensitivity; (2) acoustic-electric stimulation and acoustic-only stimulation did not systemically modulate subsequent conscious auditory perception.

**Figure 3.**
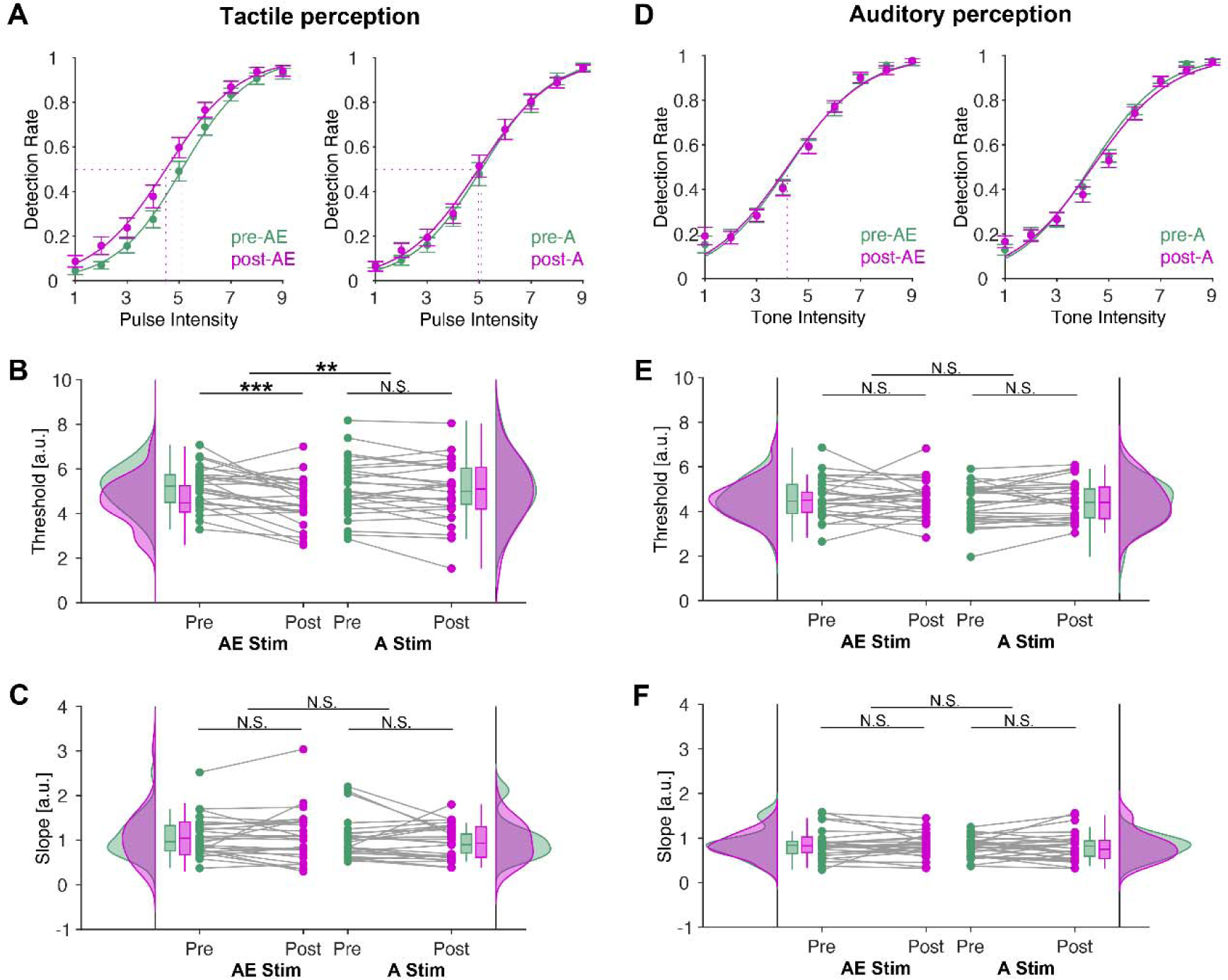
Behavioral performance in the target-detection task before and after acoustic-electric stimulation and acoustic-only stimulation. **(A).** Group-averaged psychometric function in the tactile-detection task. **(B).** The left and right pairs of plots respectively represent the tactile-detection threshold before and after acoustic-electric stimulation (left) and acoustic-only stimulation (right). The tactile threshold decreased significantly after acoustic-electric stimulation, but not after acoustic-only stimulation. **(C).** Same as (B), but for the psychometric slope. No significant effect on psychometric slope was observed after acoustic-electric stimulation or acoustic-only stimulation. **(D-F).** Same as (A-C), but for behavioral performance in the auditory-detection task. No significant effect of acoustic-electric stimulation or acoustic-only stimulation on the auditory-detection threshold or slope was observed. Data are presented as mean ± sem across participants in (A, D). The raincloud plots in (B-C, E-F) visualize the data distribution, the horizontal line within each boxplot indicates the median value across participants; the bottom and top edges of the box indicate the first and third quartile values, the whiskers indicate the most extreme values within 1.5 times the interquartile range. The dots in (B-C, E-F) represent individual participants. Green and magenta respectively represent pretest and posttest. N.S. non-significant, \*\**p* < 0.01, \*\*\**p* < 0.001.

To investigate neural responses reflecting conscious perception, we computed ERPs evoked by identical stimuli that participants detected vs. undetected. Channel-by-channel analysis revealed that detected (vs. undetected) tactile stimuli elicited a larger negativity distributed in the contralateral temporal area, which was defined as a region of interest (ROI) (Fig.S1A). The detected (vs. undetected) auditory stimuli evoked a similar negativity distributed in the frontal and central scalp areas (Fig.S1B). Detected stimuli in both modalities evoked a large positivity distributed widely over the scalp covering mainly centroparietal areas (Fig.S1). ERP waveforms averaged over the aforementioned ROIs are depicted in Fig.S2. Consistent with previous research (Al et al., 2020; Eklund and Wiens, 2019), the early negativity evoked by the detected stimuli was observed at ∼140 ms from the target onset, and a later more sustained positivity was observed 250-500 ms from the target onset (Fig.S2). These components were reliably observed across phases (pretest, posttest) and sessions. Next, an awareness-related difference waveform was obtained by subtracting the responses to undetected stimuli from the responses to detected stimuli. To assess whether acoustic-electric stimulation or acoustic-only stimulation enhanced neural responses reflecting conscious perception, we compared the awareness-related difference waveforms (detected minus undetected) after vs before stimulation. We found a significant increase in the amplitude of the late tactile component, but not the early component, after vs before acoustic-electric stimulation (late component: t_25_ = −3.875, *p* = 6.8 × 10^-4^, d = −0.760; early component: t_25_ = −0.297, *p* = 0.769, d = −0.058; Fig.4A-D). However, we found no significant change in the amplitude of either early or late tactile component after vs before acoustic-only stimulation (late component: t_25_ = −0.279, *p* = 0.782, d = −0.055; early component: t_25_ = −0.628, *p* = 0.536, d = −0.123; Fig.4A-D). Analogous to the behavioral analysis, the different effects of acoustic-electric and acoustic-only stimulation on the amplitude of the late tactile component were confirmed by a significant *stimulation* (acoustic-electric stimulation versus acoustic-only stimulation) × *time* (pretest versus posttest) interaction (F_1,25_ = 5.002, *p* = 0.034, effect size = 0.167). For conscious auditory perception, we found no significant change in the amplitude of early or late auditory component after vs before acoustic-electric stimulation (late component: t_25_ = −0.489, *p* = 0.629, d = −0.096; early component: t_25_ = 0.285, *p* = 0.778, d = 0.056) or acoustic-only stimulation (late component: t_25_ = 0.133, *p* = 0.896, d = 0.026; early component: t_25_ = −0.951, *p* = 0.351, d = −0.187) (Fig.4E-H). These results indicate that (1) acoustic-electric stimulation but not acoustic-only stimulation enhanced subsequent neural responses to conscious tactile perception and the effect applied exclusively to the late but not the early response; (2) acoustic-electric stimulation and acoustic-only stimulation did not systemically modulate subsequent neural responses to conscious auditory perception.

**Figure 4.**
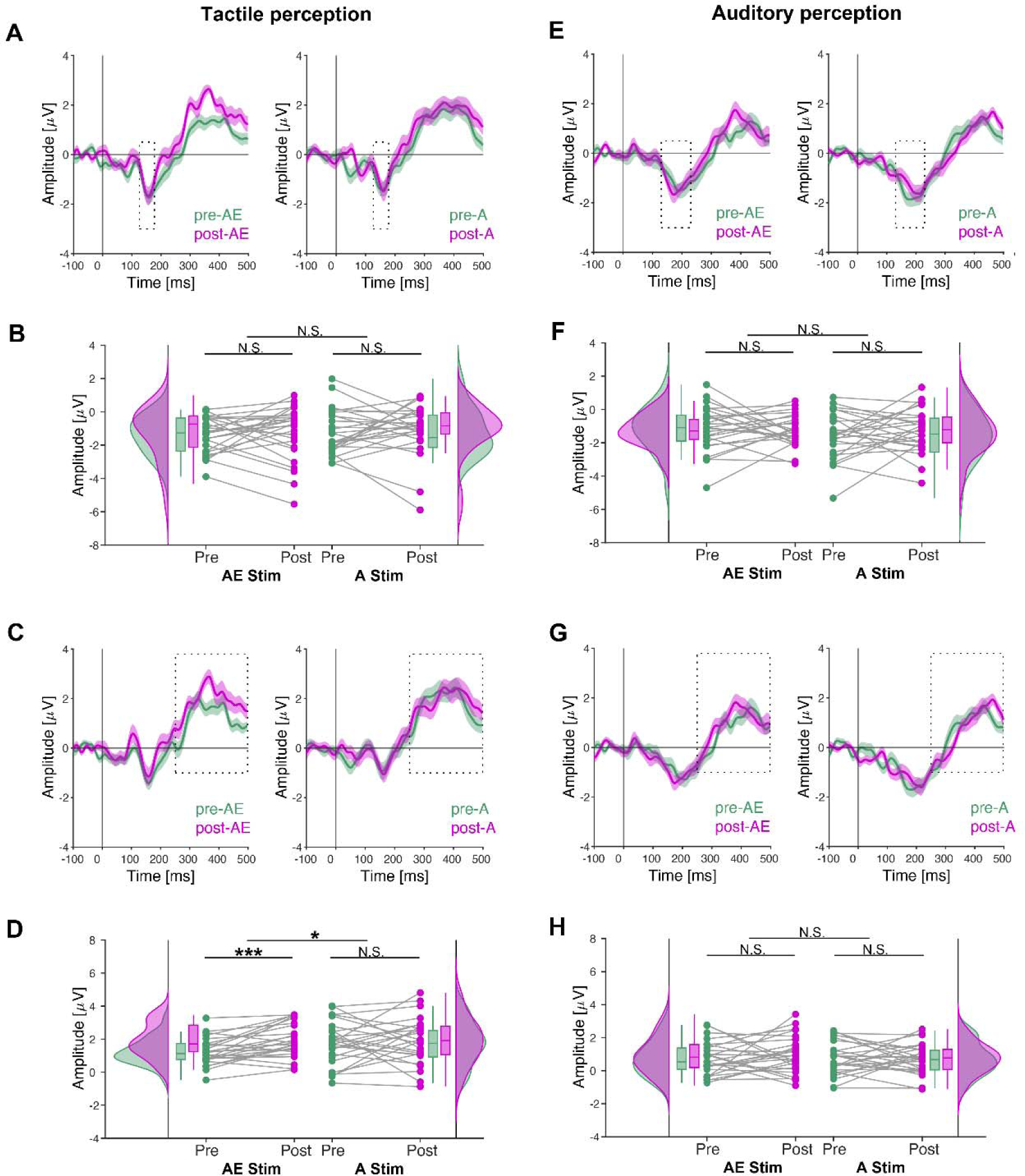
Neural responses in the target-detection task before and after acoustic-electric stimulation and acoustic-only stimulation. **(A).** Group-averaged tactile awareness-related difference waveforms of the early tactile component (averaged over ROIs). **(B).** The left and right pairs of plots respectively represent the amplitude of the early tactile component before and after acoustic-electric stimulation (left) and acoustic-only stimulation (right). No significant effect of either stimulation was observed on the early component. **(C-D).** Same as (A-B), but for the late tactile component. The amplitude of the late tactile component increased after acoustic-electric stimulation, but not after acoustic-only stimulation. **(E-H).** Same as (A-D), but for neural responses in the auditory-detection task. No significant effect of either stimulation on the early or late auditory component was observed. Data are presented as mean ± sem across participants in (A, C, E, G). The raincloud plots in (B, D, F, H) visualize the data distribution, the horizontal line within each boxplot indicates the median value across participants; the bottom and top edges of the box indicate the first and third quartile values, the whiskers indicate the most extreme values within 1.5 times the interquartile range. The dots represent individual participants. Green and magenta respectively represent pretest and posttest. N.S. non-significant, \**p* < 0.05, \*\*\**p* < 0.001.

### No significant effect of electric-only stimulation on conscious tactile perception

While the results from Experiment 1 suggest that acoustic-electric stimulation, but not acoustic-only stimulation, can improve conscious tactile perception, it remains unclear whether the effect is caused by the electric stimulation alone or its interplay with the acoustic stimulation. To address this question, we performed Experiment 2 in which we applied electric-only and electric-sham stimulation (Fig.1B, see more details in Methods) and compared tactile-detection thresholds and psychometric slopes after vs before stimulation. We found no significant change in tactile-detection threshold or slope after vs before electric-only (threshold: t_25_ = 1.490, *p* = 0.149, d = 0.292; slope: t_25_ = −0.694, *p* = 0.494, d = −0.136) or electric-sham stimulation (threshold: t_25_ = 0.179, *p* = 0.859, d = 0.035; slope: t_25_ = 1.260, *p* = 0.219, d = 0.247) (Fig.5). This result suggests that prior electric-only stimulation and/or placebo-related changes do not systemically modulate conscious tactile perception. Taken together, these results indicate that the improvement of conscious tactile perception observed in Experiment 1 was caused by the interplay of acoustic-electric stimulation, rather than by the acoustic stimulation alone (Experiment 1), the electric stimulation alone, or any placebo-related change (Experiment 2).

**Figure 5.**
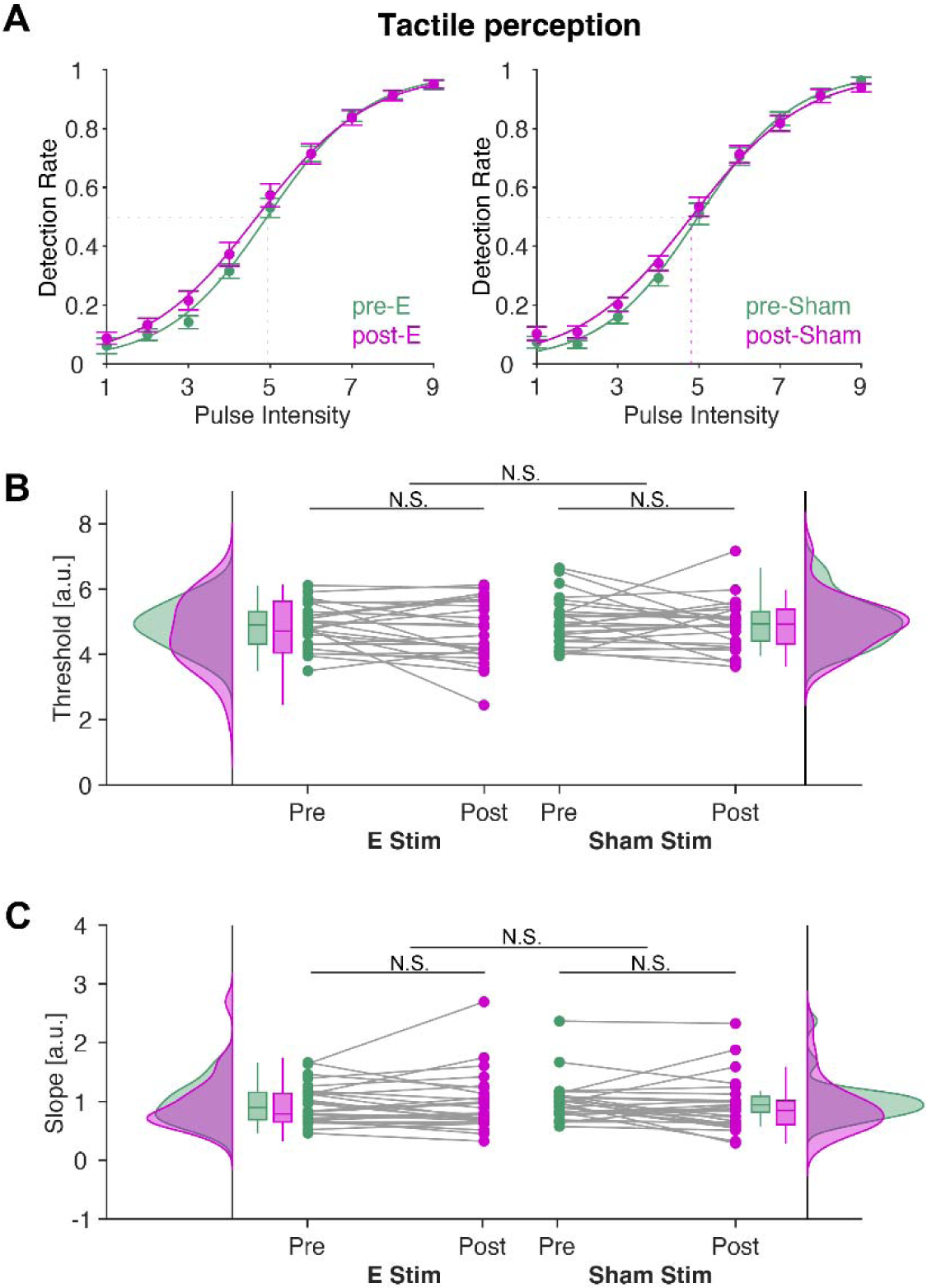
Behavioral performance in the tactile-detection task before and after electric-only stimulation and electric-sham stimulation. **(A).** Group-averaged psychometric function in the tactile-detection task. **(B).** The left and right pairs of plots respectively represent the tactile-detection threshold before and after electric-only stimulation (left) and electric-sham stimulation (right). **(C).** Same as (B), but for the psychometric slope. No significant effect of electric-only stimulation or electric-sham stimulation on tactile-detection threshold or slope was observed. Data are presented as mean ± sem across participants in (A). The raincloud plots in (B-C) visualize the data distribution, the horizontal line within each boxplot indicates the median value across participants; the bottom and top edges of the box indicate the first and third quartile values, the whiskers indicate the most extreme values within 1.5 times the interquartile range. The dots in (D-C) represent individual participants. Green and magenta respectively represent pretest and posttest. N.S. non-significant.

### Acoustic-electric stimulation modulates connections in a recurrent neural processing model

To explore potential neural mechanisms underlying the observed effects of acoustic-electric stimulation on tactile perception, we conducted a DCM analysis of tactile neural responses in the following three steps. Firstly, five models differing in the number of sources and the extrinsic connections between these sources were fitted to the tactile-evoked response (Fig.6A). Using Bayesian Model Selection (BMS), we found decisive evidence for a model in both pretest and posttest (log-model evidence for the winning model, relative to the second-best model: pretest, 3173; posttest, 2225; both corresponding to an exceedance probability of >99%) that included reciprocal connections between contralateral primary somatosensory cortex (cSI) and contralateral secondary somatosensory cortex (cSII), and between bilateral secondary somatosensory cortex (cSII, iSII) and bilateral premotor cortex (cPMC, iPMC) (Fig.6B, model 4). Secondly, six models allowing for changes in the modulatory connectivity were fitted to explain the tactile-response difference between detected and undetected stimuli (Fig.6C). A model incorporating global recurrent connections showed the greatest evidence (log-model evidence relative to the second-best model: pretest, 3223; posttest, 1570; exceedance probability >99%) and was therefore selected as the winning model (Fig.6D, model 6). This result is in accordance with the previous study that identified a role of recurrent neural processing in conscious tactile perception (Auksztulewicz and Blankenburg, 2013). Thirdly, the winning model for each subject before and after stimulation was combined into a single parametric empirical Bayes (PEB) model. Connection changes induced by the acoustic-electric stimulation were identified and calculated using Bayesian model reduction (BMR) and averaging (BMA). We found that the acoustic-electric stimulation had an excitatory effect on the connections from cSI to cSII (posterior estimate (Ep) = 0.301, a log-scaling parameter corresponding to the connection change modulated by acoustic-electric stimulation; posterior probability (Pp) = 1), from iPMC to iSII (Ep = 0.389; Pp = 1), and from iPMC to cPMC (Ep = 0.665; Pp = 1). Moreover, the acoustic-electric stimulation had an inhibitory effect on the connection from iPMC to cSII (Ep= −0.271; Pp = 1) (Fig.7A). These results indicate that the acoustic-electric stimulation modulated connection strength within a global recurrent processing model.

**Figure 6.**
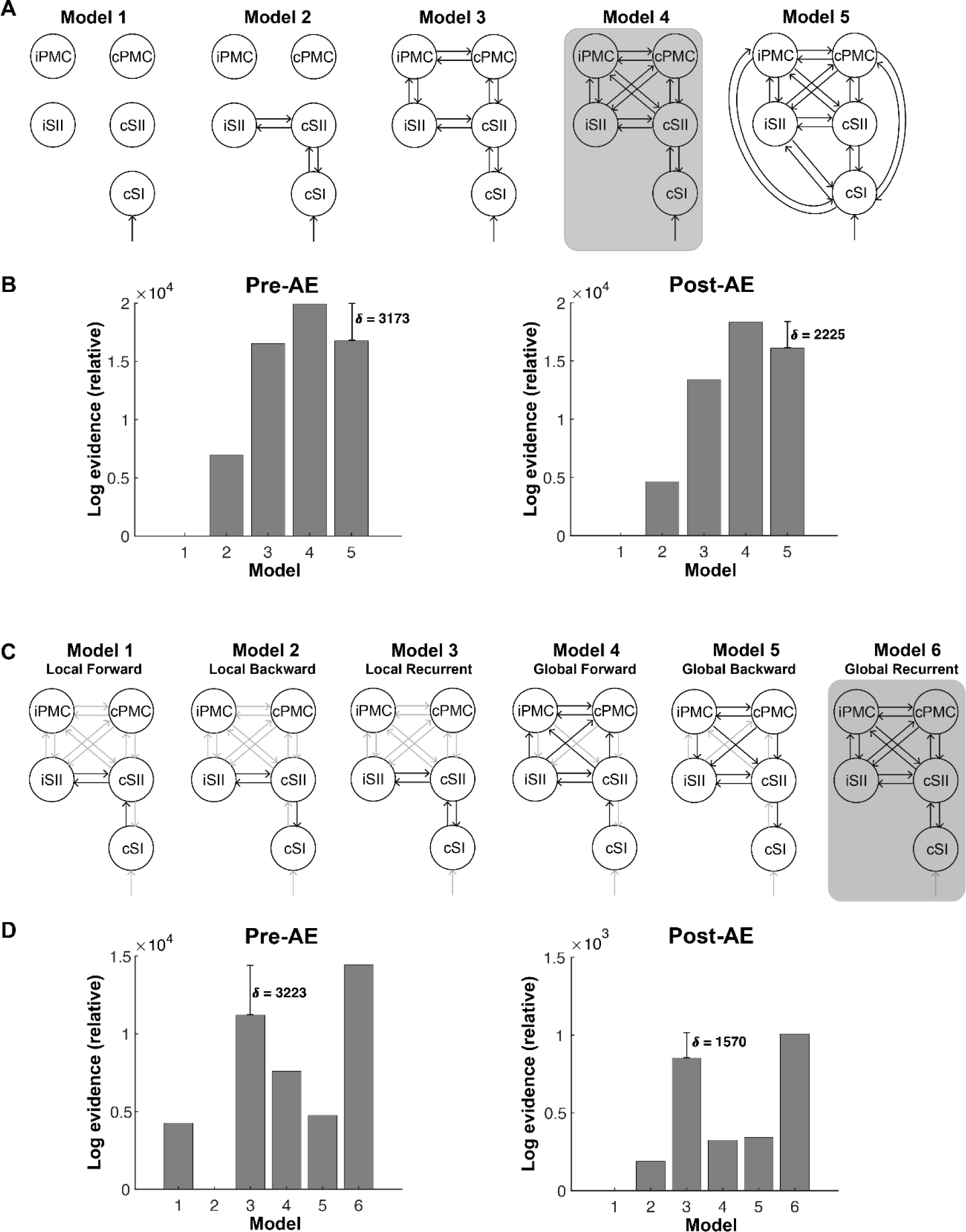
Model space of dynamic causal modeling. **(A).** Five alternative models of effective connectivity were fitted to individual participants’ ERPs evoked by the tactile stimulation. All models included the tactile input to the contralateral SI and differed with respect to the number of sources and the extrinsic connections between these sources. **(B).** Fixed-effects Bayesian model selection revealed that model 4 (shaded dark gray) outperformed the other models. Model 4 included five sources (cSI, cSII, iSII, cPMC, iPMC) and recurrent connections between cSI and cSII, and between SII and PMC. **(C).** Modeling the modulatory effect of acoustic-electric stimulation on extrinsic connections. Six models were designed in which stimulation modulated a different subset of extrinsic connections between cSI and cSII, and between bilateral SII and bilateral PMC. **(D).** Fixed-effects Bayesian model selection revealed that model 6 (i.e., global recurrent model, shaded dark gray) outperformed the other models. Model 6 allowed for recurrent connections among the five sources to be modulated by acoustic-electric stimulation.

**Figure 7.**
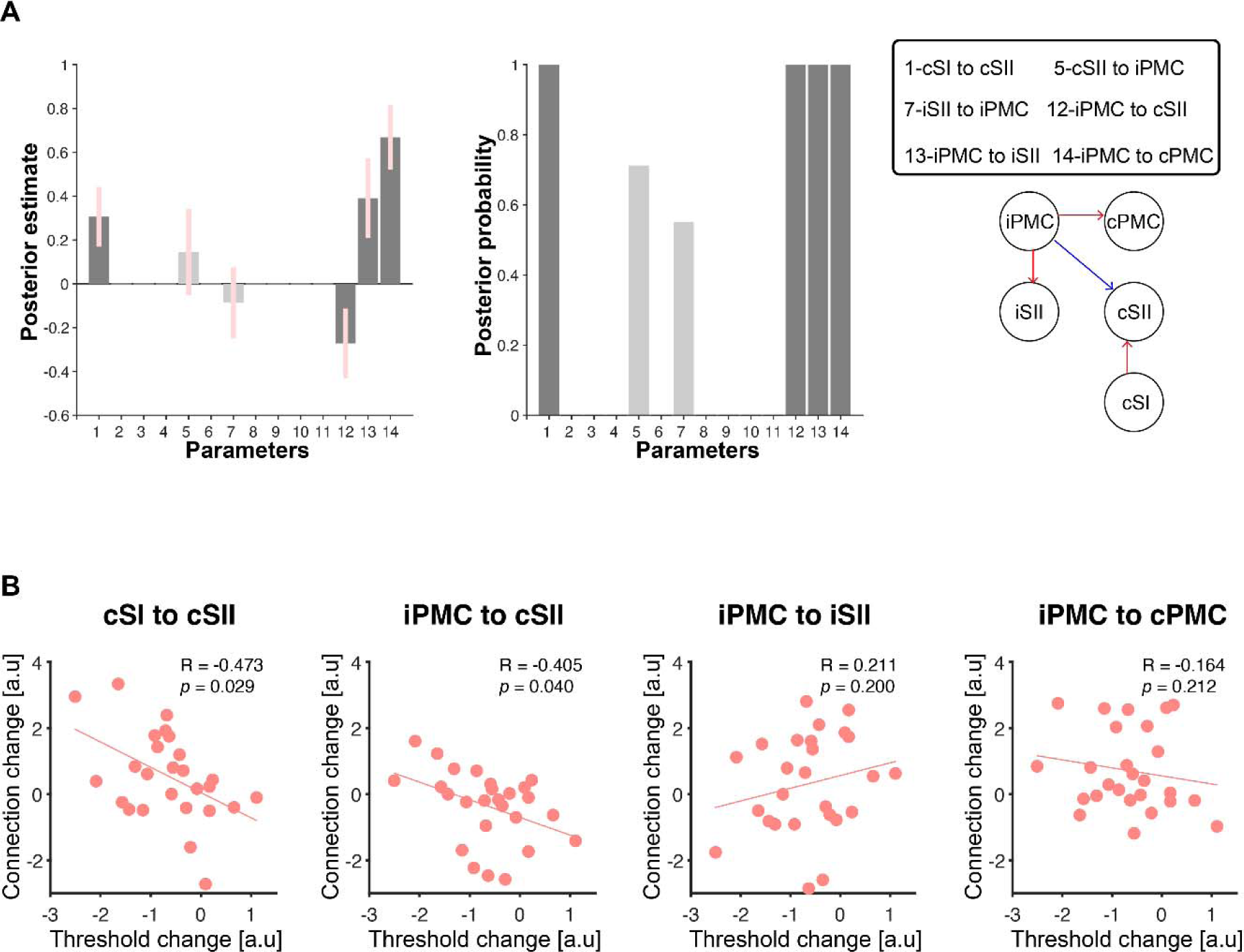
Effects of acoustic-electric stimulation on model parameters estimated using PEB. **(A).** Posterior estimates of connections (left bar plot) as log-scaling values relative to the priors and posterior probability of these connections (right bar plot). Significant changes (posterior probability > 99%) induced by acoustic-electric stimulation were observed in a subset of connections (right network graph) as the posterior estimates were significantly greater or smaller than the prior mean of zero (i.e., a log scaling of 100%): an excitation of connections between cSI and cSII, iPMC and iSII, and iPMC and cPMC (red arrows), and an inhibition of connections from iPMC to cSII (blue arrows). **(B).** The scatterplot shows results from a correlation analysis testing for an association between changes in tactile-detection threshold and changes in connectivity parameter. Correlation coefficient R and *p*-value describe, respectively, the strength and statistical significance of the coupling (linear regression line) across all participants. Supplementary Materials for

To assess whether the modulatory effects of acoustic-electric stimulation on connection were related to the observed improvements of conscious tactile perception, we next explored the correlation between connection changes (i.e., posterior estimates) and tactile-threshold changes after acoustic-electric stimulation. We found a significant negative correlation between the connection change from cSI to cSII and the change of tactile-detection threshold (R = −0.473, *p* = 0.029, false-discovery rate (FDR) corrected). We obtained a similar result for the connection change from iPMC to cSII (R = −0.405, *p* = 0.040, FDR-corrected) (Fig.7B). These results suggest that stronger improvements of conscious tactile perception were accompanied by increased bottom-up excitation from cSI to cSII and reduced top-down inhibition from iPMC to cSII.

## Discussion

We found that (1) acoustic-electric stimulation can lead to an improvement of conscious tactile perception without a concomitant change in auditory perception in healthy human participants, (2) this improvement is caused by the interplay of acoustic and electric stimulation rather than any of the unimodal stimulation alone, and (3) the effect of acoustic-electric stimulation on consciousness correlates with inter-regional connection changes in a recurrent neural processing model. Overall, our findings provide evidence that multimodal acoustic-electric stimulation can modulate conscious tactile perception and propose modulation of inter-regional connections as a potential neural mechanism.

### Acoustic-electric stimulation modulates conscious tactile perception

We found that prior acoustic-electric stimulation improved conscious tactile perception, as indicated by increases in both detection performance and the awareness-related ERP component after vs before acoustic-electric stimulation. These observations confirm and replicate previous findings in DOC patients, showing that acoustic-electric stimulation can reliably promote consciousness (Wu et al., 2022).

Gamma oscillations have been previously associated with conscious tactile perception in both observational and experimental studies (Gross et al., 2007; Meador et al., 2002; Siegle et al., 2014). For example, Siegle et al. optogenetically modulated gamma activity in mice and found improved detection of weak tactile stimulation (Siegle et al., 2014), thereby shedding light on a causal role of gamma oscillations in conscious tactile perception. Therefore, the improvements of conscious tactile perception observed in our study may be attributed to stronger gamma oscillations induced by the application of the acoustic-electric stimulation, which indeed fluctuated strongly at gamma frequency (see Methods). Although gamma oscillations were not measured in this study, it has been well-established that rhythmic stimulation can entrain rhythmic brain oscillations at the exact frequency of the applied stimulation. For example, our previous study has shown an increase of neural activity in the gamma band after multimodal acoustic-electric stimulation in the gamma band (Wu et al., 2022).

Interestingly, we found that acoustic-electric stimulation affected exclusively participants’ response criterion, not perceptual sensitivity to tactile stimulation. This finding is compatible with prior work showing effects of transcranial magnetic stimulation in gamma band on response criterion, but not perceptual sensitivity, in a visual-detection task (Chanes et al., 2013). The researchers proposed that gamma oscillations reflect sensory evidence regardless of the stimulus presence and can therefore account for response bias (Chanes et al., 2013; Riddle et al., 2019). Thus, it is conceivable that our application of gamma acoustic-electric stimulation induced gamma oscillations that reflected accumulating sensory evidence of tactile stimulation input, which then resulted in the observed shift of response criterion.

Similarly, we found acoustic-electric stimulation enhanced exclusively the late but not the early neural response to detected vs. undetected targets. The observation of significant ERP components that discriminate between detected vs. undetected stimuli in an early and a late time window is consistent with previous ERP studies on awareness (Al et al., 2020; Auksztulewicz and Blankenburg, 2013; Auksztulewicz et al., 2012; Eklund et al., 2020; Eklund and Wiens, 2019). The early negative component has been associated with perceptual sensitivity and is thought to be unaffected by response criterion (Koivisto and Grassini, 2016; Mazzi et al., 2020). As such, the lack of a significant stimulation effect on the early component in the present study is consistent with the null result on perceptual sensitivity. However, there is an ongoing controversy on whether the late component is a reliable correlate of consciousness or reflects post-perceptual processes (Cohen et al., 2020; Forster et al., 2020). Recent research has shown that the late component may not emerge in response to perceived yet unreported stimuli (stimuli in that study were irrelevant to the participants’ task) (Pitts et al., 2014). Similarly, Schroder et al. reported that the late component may not emerge from target detection when report requirements are controlled using a matching task (Schroder et al., 2021). Thus, our interpretation of the stimulation-induced increase in the late positive component as an effect on late conscious processing (as reflected by) needs to be treated with caution. It remains possible that the acoustic-electric stimulation modulated the post-perceptual reporting rather than conscious tactile perception *per se*.

### No effect of acoustic-electric stimulation on conscious auditory perception

We found that acoustic-electric stimulation did not systemically modulate subsequent conscious auditory perception. Even though a few studies have shown that transcranial alternating current stimulation (tACS) can directly modulate auditory perception in a phase-dependent manner (Neuling et al., 2012; Riecke et al., 2015), the evidence for overall or sustained benefits from tACS (compared with sham stimulation) for conscious auditory perception is scarce. For example, Riecke et al. found that the relative phase of 4-Hz tACS modulated the detection of 4-Hz click trains; however, click detection did not differ significantly under 4-Hz tACS vs. sham stimulation (Riecke et al., 2015).

A recent study revealed that the effects of gamma tACS on auditory temporal perception may depend on the difference between the stimulation frequency and the endogenous brain oscillation frequency. More specifically, tACS with frequencies 4 Hz above the endogenous gamma frequency was found to accelerate the endogenous oscillation and improve auditory gap-detection performance. In contrast, tACS with frequencies 4 Hz below the endogenous frequency did not improve auditory perception (Baltus et al., 2018). The endogenous gamma frequency has been consistently reported to exceed 45 Hz in many human studies (Baltus and Herrmann, 2015; Zaehle et al., 2010). In light of this, the gamma frequency applied in our study (40 Hz) may be lower than the typical endogenous oscillation frequency, which may explain why it did not improve auditory perception in our study. However, our interpretation requires caution, given the task differences across the studies (i.e., tone detection in our study vs. gap detection in Baltus et al.’s study).

### No effect of unimodal stimulation on conscious tactile perception

We found that prior acoustic-only or electric-only stimulation and/or potential placebo-related changes did not systemically modulate conscious tactile perception, suggesting that the observed improvement of conscious tactile perception after acoustic-electric stimulation was caused by the interplay of the acoustic and electric stimulation, rather than by the unimodal acoustic or electric stimulation alone. These findings are in line with the studies in tinnitus and AD that showed superior effects of multimodal stimulation over unimodal stimulation (see Introduction).

One potential explanation could be that the unimodal acoustic and electric stimulation have simple additive effects. However, this interpretation is difficult to reconcile with our observation of non-significant effects of either unimodal acoustic stimulation or unimodal electric stimulation, rendering this alternative less plausible. Another, perhaps more intriguing explanation is that the consciousness benefits were driven by the integration of the acoustic and electric inputs. Indeed, this notion has been supported by animal studies. For example, auditory-tactile integration has been found in the primary auditory cortex of monkeys (Lakatos et al., 2007). Moreover, animal studies in mice have shown that auditory stimulation and forepaw tactile stimulation activate neurons in the somatosensory cortex. When the tactile stimulation was paired with the auditory stimulation, the neuronal activation in somatosensory cortex and behavioral performance in a tactile-based task were significantly improved. These results indicate that the somatosensory cortex can encode multisensory information and auditory input to it can enhance its sensory encoding, leading to changes in responses to tactile input (Godenzini et al., 2021). Although we could not directly extract participants’ somatosensory cortical activity, our DCM results suggest a neuronal state change in the somatosensory cortex after acoustic-electric stimulation, which we discuss further below.

Our null results based on unimodal electric stimulation are consistent with a recent tACS study that applied tACS over SI at alpha, beta, and gamma frequencies and found no significant change in near-threshold tactile perception and tactile temporal discrimination threshold (Manzo et al., 2020). However, it remains possible that unimodal stimulation did not reach the threshold for promoting consciousness. Future research may examine this possibility by e.g. applying the stimulation at an increased intensity.

### Acoustic-electric stimulation modulates connections in a recurrent neural processing model

Our finding that a recurrent neural processing model can account for cortical responses reflecting conscious tactile perception aligns well with results from a prior study (Auksztulewicz and Blankenburg, 2013). We found that acoustic-electric stimulation had a modulatory effect on the following inter-regional connections within the recurrent processing model: from cSI to cSII, from iPMC to iSII, and from iPMC to cPMC, and from iPMC to cSII. These findings suggest that the acoustic-electric stimulation affected both somatosensory areas and PMC, and modulated the strength of the direct connections between them. However, it remains possible that the observed alterations in inter-regional cortical connections may be driven indirectly by more unspecific mechanisms, such as changes in arousal or alertness, rather than by the acoustic-electric stimulation directly. Furthermore, we observed that the connection changes from cSI and iPMC to cSII were negatively correlated to the stimulation effects on tactile perception. This suggests that the observed improvement of conscious tactile perception may be accompanied by greater excitatory projection from cSI to cSII and weaker inhibitory projection from iPMC to cSII. The alterations of these two putative connections to cSII might point to an essential role of SII in conscious tactile perception. According to previous studies, the SI and SII are responsible for transforming tactile input into tactile perception, and the SII is the first cortical region to show response differences between detected and undetected tactile stimulation (Auksztulewicz et al., 2012; Schroder et al., 2019; Wuhle et al., 2010). In combination with our modeling results, these results converge on the idea that SII is a crucial region for conscious tactile perception.

In conclusion, our study provides evidence that multimodal stimulation combining music stimulation and electrical TNS can promote conscious tactile perception in healthy human participants and suggests modulation of inter-regional cortical connections as a plausible mechanism. No such perceptual benefit could be achieved by unimodal acoustic and electric stimulation alone. In sum, this study provides insight into the reliability and functional principle of multimodal acoustic-electric stimulation, which can inspire the application of this novel approach in clinics to promote patients’ consciousness.

## Materials and Methods

### Participants

Participants were recruited from the student population of Maastricht University. Fifty-four participants (42 females, 12 males; ages: 18-30 years) completed the study. Two participants with poor EEG data quality (Experiment 1, see section *Electrophysiology* for details) were excluded. The remaining 52 participants were included for further analysis: 26 participants in Experiment 1 and another 26 participants in Experiment 2. All participants reported normal hearing and no history of neurological or psychiatric disorders. No participants had contraindications to transcutaneous electric stimulation. Written informed consent was obtained prior to the experiment. Participants were compensated with study credits or monetary reward for their participation. The experimental procedure was approved by the local research ethics committee (Ethical Review Committee Psychology and Neuroscience, Maastricht University).

### Study overview

The study included two double-blinded, randomized, crossover experiments. Each experiment consisted of two sessions separated by at least 48 hours. Each session comprised three phases: an assessment of conscious perception before stimulation (i.e., pretest), application of the stimulation, and an assessment of conscious perception after stimulation (i.e., posttest) (Fig.1). The only difference between two sessions was the type of applied stimulation: in Experiment 1, it was either acoustic-electric or acoustic-only, and in Experiment 2, it was either electric-only or electric-sham. The order of stimulation type was counterbalanced across participants. Participants and data collectors were blinded to stimulation type throughout the experiment. During the pretest and posttest phases, behavioural responses (Experiments 1 and 2) and neural responses (Experiment 1) were measured in tactile (Experiments 1 and 2) and auditory (Experiment 1) detection tasks.

## Experiment 1

### Conscious perception assessment

To allow assessing conscious perception during the pretest and posttest phases, a target-detection task adopted from previous studies was used (Auksztulewicz and Blankenburg, 2013; Eklund and Wiens, 2019; Sanchez et al., 2020; Schroder et al., 2021). Tactile stimuli were 1-ms biphasic square wave pulses generated by a constant current stimulator (DS5, Digitimer). The tactile stimuli were delivered via Ag/AgCl electrodes adhered to the tip of the left index finger to stimulate the median nerve. Auditory stimuli were 1000-Hz pure tones with a duration of 100 ms (including 5 ms fade-in and 5 ms fade-out), embedded in continuous white noise (44.8 dB SPL). The auditory stimuli were presented binaurally via insert earphones.

Prior to the main assessment, individual tactile and auditory target-detection thresholds were measured with a two-step procedure involving the method of adjustment (step 1) and the method of constant stimuli (step 2). In step 1, participants were asked to increase the current intensity from 0.3 mA (in steps of 0.1 mA) to the lowest stimulus intensity at which they could detect the electric pulse. In step 2, participants were exposed to pulses with various intensities (ten equidistant levels centred on the current intensity determined in the first step, with an increment size 0.06 mA). A total of 100 trials (10 for each intensity) were presented in random order. Participants responded with a button response on each trial to indicate whether they had detected the pulse or not. The obtained data were fitted with a logistic psychometric function, from which three intensities yielding the following performance levels were derived: 1% detection rate (intensity 1), 50% detection rate (intensity 5, equal to detection threshold), and 99% detection rate (intensity 9). Intensities 1 and 9 were used to define the intensity range that was presented in the subsequent assessment of conscious perception. The auditory-threshold assessment followed the same procedure as the tactile-threshold assessment above, with the exceptions that the loudness of the tone started at an audible level (45 dB) and decreased in steps of 4 dB in the first step, and the increment was 2 dB in the second step.

After the threshold measurements, four blocks of detection-task trials were presented while EEG was recorded. Each block contained 150 trials and lasted ∼9 min. Two blocks contained only tactile trials and two blocks contained only auditory trials. The presentation order was TATA or ATAT (T: tactile; A: auditory) and counterbalanced across participants. In each block, the 150 trials were presented at nine intensities, which were equidistantly spaced between 1% (intensity 1) and 99% (intensity 9) detection rate, as described above. The number of trials presented at each intensity level followed a normal distribution to maximize the trials with intensities near the detection threshold (Fig.2B).

Each block started with the presentation of a task instruction on the screen, which instructed participants to perform the tactile or auditory task. Each trial started with a white central fixation cross on a black screen for 2000 ms. A tactile or auditory stimulus was applied with a random delay between 1000-1500 ms after the fixation onset. Next, a response screen was displayed, instructing participants to report whether they had detected a stimulus or not by pressing one of two buttons (“1” or “2”) within 1500 ms. No feedback on task performance was given. Buttons “1” and “2” in first two blocks represented “Yes: detected” and “No: undetected”. To control for motor-response mapping, the mapping was reversed in blocks 3 and 4 with “1” and “2” indicating “No: undetected” and “Yes: detected” respectively (Fig.2A). Participants could take a break after each block for as long as they needed. Two brief practice blocks of 30 trials were presented prior to the main assessment to familiarize participants with the stimuli and tasks. The threshold measurement and practice procedures were conducted only in the pretest phase of each session.

### Acoustic and electric stimulation

Acoustic stimulation consisted of eight pieces of Japanese music. To control for a potential learning effect, the eight pieces of music were divided into two sets that were played in individually randomized order in the two sessions. The onset/offset of each excerpt was ramped up/down using 5-s long ramps. Excerpts were amplitude-compressed (compression ratio: 120:1, threshold: −12 dB) and sequenced to form a continuous 20-min stream of music. To enforce rhythmic brain activity at gamma frequency, the sequence was amplitude-modulated at a frequency of 40 Hz (sinusoidal modulation, depth: 100%). The amplitude of the overall sequence was scaled to avoid clipping. The acoustic stimulation was presented diotically through insert earphones at a comfortable sound level (70.8 dB SPL).

Electric stimulation consisted of the non-invasive application of alternating currents to the participant’s face to facilitate rhythmic trigeminal nerve activity. Analogously to the acoustic stimulation, the current waveform was a sinusoid with a frequency of 40 Hz. The current was applied using two pairs of square-shaped rubber electrodes (size: 3 cm × 3 cm) placed at the bilateral middle and lower part of the participant’s face to stimulate the second and third branches of the trigeminal nerve (i.e., the maxillary nerve and the mandibular nerve, respectively). Based on previous research and pilot tests (McGough et al., 2019), the intensity of the current was fixed to ±4 mA. The onset/offset of the current was ramped up/down using 5-s long ramps. The electrodes were adhered to the participant’s skin using conductive paste and the impedance was kept below 10 kΩ. The electric stimulation was delivered using battery-powered DC stimulators (NeuroConn, Germany).

Acoustic and electric stimuli were digitally generated using a sampling rate of 16 kHz and then converted simultaneously to analog signals using a multi-channel D/A converter. In Experiment 1, phase-locked acoustic stimulation and electric stimulation were simultaneously presented for 20 min during the acoustic-electric stimulation period. During the acoustic-only stimulation period, only acoustic stimulation was presented for 20 min but the electric stimulation was slowly ramped down after the initial 30 s. The continuous presentation of acoustic stimulation ensured that participants did not receive auditory cues differentiating the two stimulation conditions. The 30-s electric stimulation at the beginning induced sensations similar to the electric stimulation in the main stimulation condition. Together these measures served to blind participants to the stimulation conditions.

Upon completion of each session, participants were asked to complete a questionnaire in which they were asked to (1) guess the time course of the delivered electric stimulation (“no stimulation”, “beginning”, “end”, “continuous”), (2) report if they suffered any side effects, (3) report their level of familiarity with the presented music.

## Experiment 2

The assessment of conscious perception in Experiment 2 was similar to that in Experiment 1, with the exceptions that only two tactile blocks were presented and only behavioral data were recorded. The stimulation administered in Experiment 2 was also similar to that in Experiment 1, except that the acoustic stimulation was removed, resulting in electric-only and electric-sham stimulation (Fig.1B).

## Data acquisition and processing

### Behavior

Trials without responses were rejected and the remaining trials were classified as “detected” or “undetected” depending on the participant’s response. The detection rates were calculated as a function of stimulus intensity (from 1 to 9) and then fitted by a logistic function using a maximum likelihood criterion, as implemented in the Palamedes toolbox(Prins and Kingdom, 2018). The function is given by:

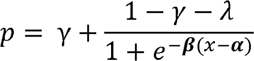

where *γ* and *λ* are the lower and upper bounds of the psychometric function, reflecting the guessing rate and lapse rate; α is a threshold parameter indicating the center of the psychometric function’s dynamic range; β is related to the slope of the function. The guessing rate and lapse rate were fixed at zero and the other two parameters, detection threshold and slope, were set as free parameters. In this study, the threshold is the stimulation intensity yielding 50% detection rate on the fitted psychometric function and the slope is the steepness of the function at that intensity. The psychometric function was fitted separately for each participant, phase (pretest and posttest), session, and sensory task modality (tactile and auditory).

### EEG

#### Recording

EEG signals were recorded during the target-detection task in Experiment 1 using 39 scalp EEG electrodes (BrainCap, Brain Products LiveAmp 64, Gilching, Germany) placed according to a standard 10–20 system. The AFz electrode was used as the ground electrode and the left mastoid electrode (A1) was used as an online reference electrode. Vertical electrooculogram was recorded by placing an extra electrode below the left eye. An additional electrocardiogram electrode was placed under the left breast to record heart activity (not analysed in the present study). Electrode impedances were kept below 10 kΩ throughout the experiment. EEG recordings were online bandpass-filtered between 0.01 and 70 Hz, and digitized with a sampling rate of 500 Hz.

#### Data pre-processing and ERP analysis

Data preprocessing and analysis was performed using EEGLAB 2019.1 (Delorme and Makeig, 2004) and MATLAB 9.4. First, bad channels with a leptokurtic voltage distribution (i.e., kurtosis higher than five) were replaced by interpolating between the surrounding electrodes (spherical spline interpolation; mean number of interpolated channels across participants: 1.0). Then, the interpolated channel data were re-referenced to the average of left and right mastoids and were band-pass filtered between 0.5 Hz and 30 Hz using a Butterworth Infinite Impulse Response (IIR) filter (zero phase shift, filter order: 6). Next, independent component analysis was performed, and artifactual components were identified and discarded (mean percentage of artifactual components: 15.4%) using the EEGLAB plugin ICLables (Pion-Tonachini et al., 2019). The continuous data were segmented into epochs from −100 to 500 ms relative to stimulus onset and artifactual epochs with amplitudes exceeding ±65 μV were removed (mean percentage of excluded epochs: 0.8%). One participant’s data were excluded from further data analysis because of excessive artifacts resulting in 11.7% of epochs removed. The artifact-free, epoched data were corrected for baseline drifts by subtracting the average amplitude from −100 to −10 ms relative to the stimulus onset from the epoch.

To control for stimulation-related confounds (i.e., number of epochs, stimulation intensity) in the analysis, we balanced the number of epochs between detected and undetected conditions per intensity level using subsampling in the condition with a larger number of epochs. Compared to randomly selecting an equal number of epochs, this method was designed to minimize the interval difference (i.e., the presentation interval) between the conditions (i.e., detected vs. undetected), and control for the time-related confounds (e.g., fatigue and adaptation). One participant was excluded from further data analysis due to a limited number of epochs left (13.3%) after balancing. Overall, the average number of retained tactile and auditory epochs was 118 and 140, respectively, with the greatest number of epochs near the detection threshold as expected (Fig.2C). Finally, an equal number of detected and undetected epochs were averaged in the time domain, creating time-locked ERPs for each electrode, each condition (detected and undetected), each modality (tactile and auditory) and each participant.

ERP components (and corresponding time windows for analysis) were chosen based on previous research, with particular focus on an early negative component (tactile: 125-180ms; auditory: 120-200ms) and a late positive component (250-500ms) related to conscious perception (Dembski et al., 2021). Mean amplitudes were calculated using time-window averages at each channel, separately for detected and undetected trials. To find the channels showing significant differences related to conscious perception, the amplitudes for detected and undetected trials were compared per channel with paired t-tests and the FDR was controlled to correct for multiple comparisons. Channel clusters that were found to show significant differences between detected and undetected trials in each phase and each session were defined as ROIs. Finally, difference (detected minus undetected) waves were computed for each phase and each session, and the amplitudes of the early and late ERP components were averaged across the aforementioned time windows and the ROIs.

## Dynamic causal modeling

DCM is an effective connectivity analysis in which sensor-level data (here: ERPs) are fitted with a generative model of observed responses (David et al., 2005). The model consists of two main components: a neuronal model, describing dynamics of neural activity in a network comprising effective (directional) connectivity parameters between network nodes (brain regions), and an observation model, mapping hidden states (neural activity estimates) to sensor-level measurements. Models are fitted to the data using variational Bayesian methods (Kiebel et al., 2009) to obtain estimates of log-model evidence (enabling a comparison between alternative models) and model parameters (e.g., connectivity estimates). The connectivity parameters comprise baseline estimates shared across experimental conditions (“A matrix”), modulatory parameters denoting connectivity changes due to experimental conditions (“B matrix”; e.g., stimulus detection), and input parameters (“C matrix”, defining network nodes receiving sensory inputs). A detailed formulation of DCM for ERPs and its application to tactile detection can be found in a previous study (Auksztulewicz and Blankenburg, 2013). In the present study, DCMs were used to identify (1) a neural network model explaining ERP differences to detected and undetected stimuli (i.e., modeling conscious perception effects) and (2) connection changes in the model induced by the acoustic-electric stimulation. This analysis was conducted using the SPM 12 toolbox (http://www.fil.ion.ucl.ac.uk/spm/).

The DCMs were fit to the stimulus-evoked ERPs from 1 to 500 ms peristimulus time using an ERP convolution-based neural mass model (Moran et al., 2013). Based on findings from a related study using DCM (Auksztulewicz and Blankenburg, 2013), cSI, cSII, iSII, cPMC and iPMC were defined as sources constituting network nodes. The locations (MNI coordinates) of the five sources were obtained from previous studies and were as follows: cSI (52, −22, 44), cSII (52, −16, 16), iSII (−58, −20, 16), cPMC (30, −20, 54), iPMC (−30, −20, 54)(Auksztulewicz and Blankenburg, 2013; Wacker et al., 2011).

Firstly, the basic DCM architecture (“A matrix”) was identified using the ERPs evoked by both detected and undetected tactile stimuli. Five alternative models were constructed differing in the number of sources and the extrinsic connections between them (Fig.6A). Each model was fitted separately for each participant, resulting in 5×26 models in pretest and posttest. These models were then compared using fixed-effects BMS, which resulted in the selection of a winning model that fitted the data best across all participants (Stephan et al., 2010).

Secondly, the selected winning model was optimized with respect to the condition-dependent changes (“B matrix”) in extrinsic connections to explain the observed tactile ERP difference between detected and detected trials. Six models were constructed differing with respect to their extrinsic connections (Fig.6C). Three models assumed modulatory connections between cSI and cSII/iSII and the other three models added modulatory connections between SII and PMC. The exogenous tactile information is modeled as a direct input entering cSI (“C matrix”). As above, each model was fitted separately for each participant, and BMS was used to select the winning model.

To estimate the changes of model parameters caused by acoustic-electric stimulation, a second-level analysis was conducted using PEB (Rosch et al., 2019; Zeidman et al., 2019). PEB is used to estimate commonalities and differences in model parameters at the group level, modelled as a second-level design matrix (typically, under the assumption that parameters are normally distributed in the participant sample). Our PEB design matrix contained two columns: the first column represented the group mean (i.e., 1), and the second column represented time (i.e., −1 for pretest, 1 for posttest). The winning model from the previous step was entered per participant and time into the PEB, and subjected to BMR and BMA to prune away model parameters that did not contribute to the model evidence. To enable statistical inference, we retained model parameters at a posterior probability higher than 0.99 (i.e., strong evidence for a change of the parameters induced by the acoustic-electric stimulation).

## Statistical analysis

To test for successful participant blinding, Fisher’s exact test was run to determine if the perceived stimulation type differed between two stimulation sessions in both experiments. To assess the effect of stimulation on behavioral and neural measures within each stimulation session, two-sided paired t-tests were used. To test whether the effect differed between stimulation sessions, a two-way repeated-measures ANOVA with two within-subjects factors “*stimulation*” (acoustic-electric vs. acoustic-only in Experiment 1; electric-only vs. electric-sham in Experiment 2) and “*time*” (pretest vs. posttest) was used. The correlation between behavioral and neural effects of stimulation and its significance were assessed using Pearson’s correlation coefficient *R*. A significance criterion α = 0.05 was used and type-I error probabilities inflated by multiple comparisons were corrected by controlling the FDR.

## Acknowledgments

We thank S. Kuang for the assistance in data collection. We thank Y.X. Feng for creating the head models in Fig. 1. This work was supported by Maastricht University and China Scholarship Council (CSC 201906320078 to M.W.).

## Author contributions

Conceptualization: M.W., and L.R. Methodology: M.W., A.R., and L.R. Investigation: M.W. Visualization: M.W. Supervision: A.R., and L.R. Writing—original draft: M.W., and L.R. Writing—review & editing: A.R., and L.R.

## Data and materials availability

Data will be available in Open Science Framework (https://osf.io/y9n84/) upon publication.

---

**This PDF file includes:**

Figs. S1 to S2

**Figure S1.**
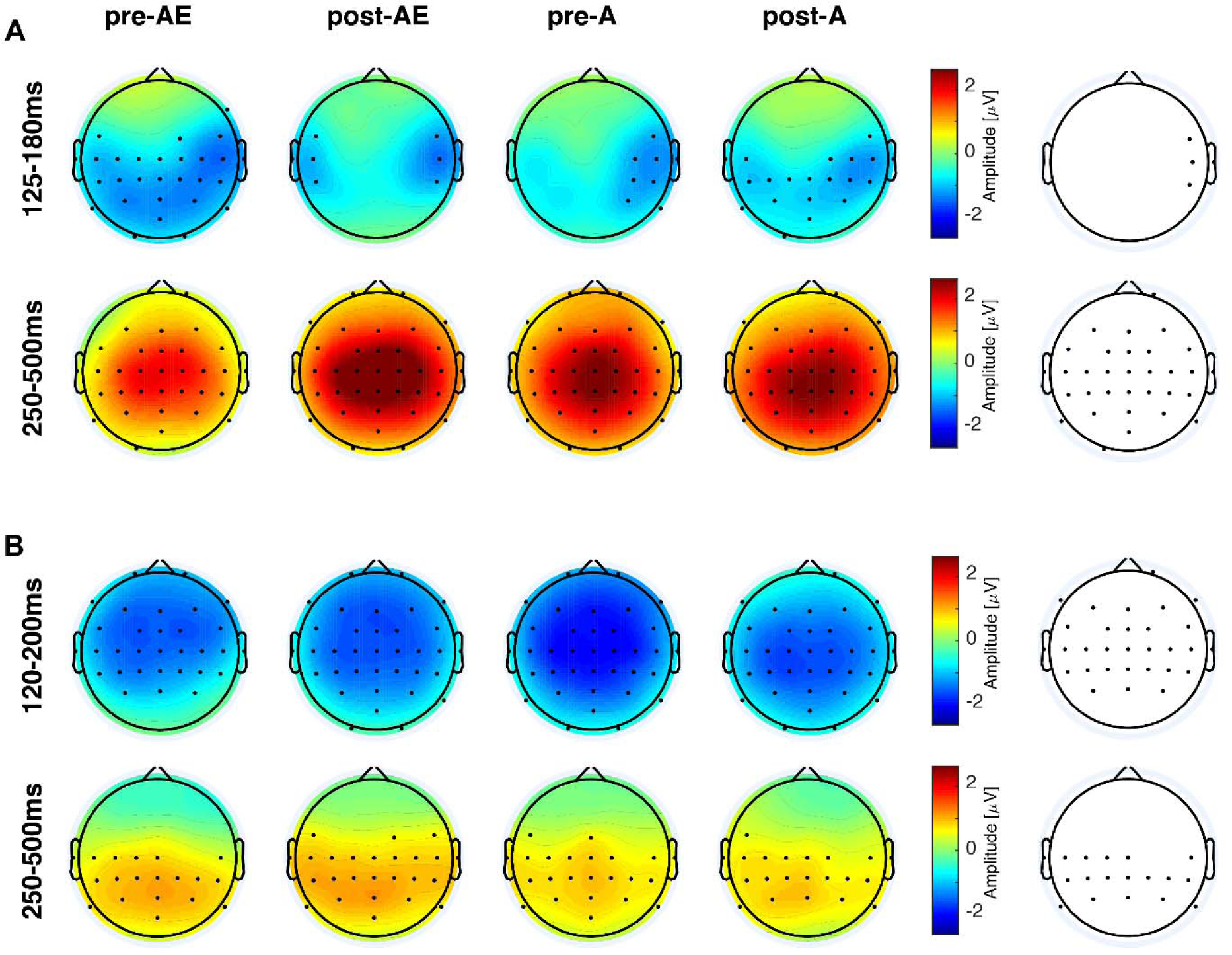
Topographic maps of the amplitude of awareness-related components (detected minus undetected) before and after acoustic-electric stimulation and acoustic-only stimulation. **(A).** The early tactile negative component was distributed over a right temporal area (upper), while the late tactile positive component was distributed over a central parietal area (bottom). **(B).** The early auditory negative component was distributed over frontal and central areas (upper), while the late auditory positive component was distributed over a parietal area (bottom). Black dots in the colored maps indicate the electrodes showing a significant difference in ERP amplitude between detected and undetected stimuli (*p* < 0.05, FDR corrected). Channels that showed such awareness-related effects reliably across all phases (pretest and posttest) and all sessions are shown in the rightmost plots.

**Figure S2.**
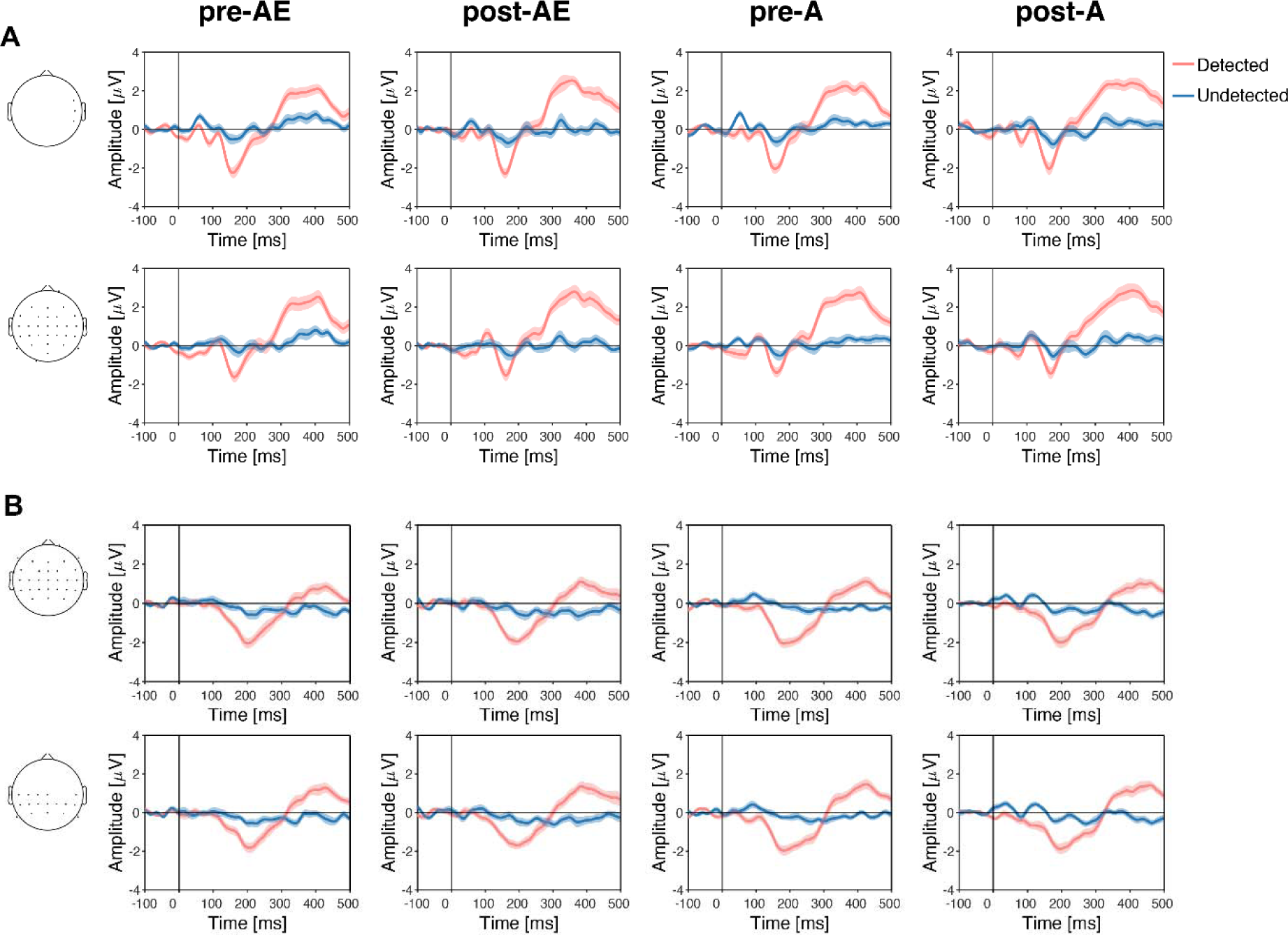
Neural responses to detected and undetected stimuli before and after acoustic-electric stimulation and acoustic-only stimulation. **(A).** Group-averaged ERP waveforms to detected (red line) and undetected (blue line) tactile stimuli averaged over a right temporal area (i.e., ROIs of the early tactile component) and a central parietal area (i.e., ROIs of the late tactile component). An early negativity and a late positivity were elicited by detected stimuli but not undetected stimuli. **(B).** Same as (A), but for the responses to auditory stimuli. ROIs are depicted with black dots on a head icon on the left (same as Fig.S1, right). Data are presented as mean ± sem across participants.

